# RNA-Binding Protein NF90 Mediates Polycomb-Independent Transactivation by EZH2 to Promote Cancer Growth

**DOI:** 10.64898/2025.12.02.691892

**Authors:** Yuan Wang, Liu Peng, Xiaodong Lu, Hongshun Shi, Mohan Zheng, Sambhavi Senthil, Jawad Akhtar, Yongik Lee, Hana Chandonnet, Jonathan D. Licht, Ximing Yang, Jonathan C. Zhao, Jindan Yu

## Abstract

Increasing evidence suggests critical roles of the polycomb-independent transactivation function of EZH2 in promoting some cancers, such as prostate cancer (PCa), yet the underlying mechanism remains poorly understood. Here, we identify the RNA-binding protein NF90 as a key mediator of this activity. NF90 interacts with EZH2, but not with other core components of the polycomb repressive complex 2 (PRC2), through its RNA-binding modules. Conversely, EZH2 engages NF90 via its intrinsically disordered RNA-binding domain in an RNA-dependent manner. NF90 and EZH2 mutually recruit each other to the AR promoter, where they cooperatively activate AR transcription and enhance downstream AR signaling. This NF90-EZH2 complex is essential for PCa cell growth: depletion of either factor abolishes proliferation, an effect rescued by AR re-expression. Similar to EZH2, NF90 promotes cell-cycle gene expression, is upregulated in advanced PCa, and is associated with poor clinical outcomes. Collectively, our findings uncover RNA-mediated protein interactions as a central mechanism underlying PRC2-independent transcriptional activation by EZH2 and establish NF90 as a major EZH2 coactivator, a master regulator of the cell cycle, and a promising therapeutic target in advanced PCa.

## INTRODUCTION

Enhancer of Zeste Homolog 2 (EZH2) is the catalytic subunit of the Polycomb Repressive Complex 2 (PRC2), wherein it interacts with other core components EED and SUZ12 to catalyze trimethylation of histone H3 at lysine 27 (H3K27me3)^1^. The resulting trimethylation facilitates chromatin condensation and target gene silencing, which are critical for PRC2’s role in development^2–4^. EZH2 dysregulation, either overexpression^5,6^ or gain-of-function mutations (e.g., Y646)^7,8^, has been linked to a wide range of cancers and shown to promote tumorigenesis. For instance, high EZH2 levels have been correlated with worse prognosis in prostate cancer (PCa), and depletion of EZH2 markedly inhibits PCa cell proliferation and survival^5,9^. Through its canonical, methyltransferase-dependent activity, EZH2 suppresses the expression of key tumor suppressor genes, including NOV^10^, and SLIT2^11^, promoting tumor progression. Additionally, EZH2 has been shown to promote tumorigenesis by methylating non-histone substrates such as FOXA1^12^, RORα^13^, GATA4^14^, and STAT3^15^, thereby modulating their stability and/or activities. In recent years, several EZH2 methyltransferase inhibitors have entered various stages of drug development and clinical trial^16–18^. Unfortunately, these drugs have shown very limited therapeutic efficacy as a single agent^19–21^. This may be in part due to non-catalytic functions of EZH2 in tumorigenesis, including recruiting non-PRC2 cofactors to promote transcriptional activation^9,22–24^. Our previous work demonstrated that EZH2 directly induces transcription of the androgen receptor (AR) gene in a Polycomb- and methylation-independent manner, resulting in PCa resistance to EZH2 enzymatic inhibitors^24^. Other studies revealed that EZH2 interacts with co-activators such as histone acetyltransferase p300 to induce transcription and promote tumor cell growth^9,23,25–27^. However, the mechanism by which EZH2 is recruited to its activating target genes, such as AR, has remained an unclear critical question.

Nuclear Factor 90 (NF90), encoded by the Interleukin Enhancer Binding Factor 3 (*ILF3*) gene, is a double-stranded RNA-binding protein (dsRBP)^28–31^. *ILF3* gene produces two major isoforms, NF90, and its longer variant NF110 that uniquely possesses a C-terminal GQSY-rich region that is absent in NF90^30^. NF90/NF110 forms a complex with nuclear factor 45 (NF45), which is important for their stability and activity^32^. The complex binds mRNAs through its tandem double-stranded RNA-binding motifs (dsRBMs), recognizing structured elements within the 5′ and 3′ UTRs to regulate mRNA stability and translation^33–35^. Moreover, NF90 also interacts with noncoding RNAs (ncRNA) through its dsRBMs and C-terminal RGG domain, recognizing structured elements such as hairpins and stem-loops^33–37^. Through these interactions, NF90 regulates RNA processing, stability, translation, and subcellular localization^36–39^. Most NF90 resides in the nucleus, tethered by RNA bound to its dsRBM^29^. In recent years, NF90 dysregulation has been reported in several malignancies, including ovarian^40^, liver^41^, cervical^42^, bladder^43^, and esophageal squamous cell carcinoma (ESCC)^44^, regulating tumorigenesis primarily through its RNA-binding activity. NF90/NF110 was also shown to participate in transcriptional regulation^30,45,46^, and work with transcriptional co-factors such as eIF2^47^ and ku proteins^48^. ChIP-seq of NF90/NF110 in erythroleukemia cells revealed that NF90 associates with chromatin and preferentially activates pro-proliferation transcription factors, thereby promoting tumor growth^49^. However, this important NF90/NF110-NF45 complex has not been studied in the context of PCa.

Here, we identify NF90 as an interacting partner of solo-EZH2, which functions outside of the PRC2 complex. NF90 and EZH2 physically interact with each other through their RNA-binding domains in an RNA-dependent manner. This RNA-mediated association enables them to mutually recruit each other to the AR promoter, thereby inducing AR expression and enhancing downstream AR signaling. Depletion of either NF90 or EZH2 markedly reduces AR expression and inhibits PCa cell proliferation, which is rescued by AR re-expression. Like EZH2, NF90 is significantly upregulated in high-grade PCa and castration-resistant PCa (CRPC), and is a major promoter of the cell cycle. In summary, our study demonstrates that RNA binding is essential for EZH2’s function as a transcriptional activator and identifies NF90 as a key co-activator of solo-EZH2, collaboratively inducing AR expression and cell growth.

## RESULTS

### The NF90/NF110-NF45 complex interacts with solo EZH2 independently of PRC2

To identify novel transcriptional activators that interact with EZH2, we examined immunoprecipitation-mass spectrometry (IP-MS) data^50^ and identified NF90 as a potential interacting partner of EZH2. To validate this interaction, we first overexpressed haemagglutinin (HA-tagged) EZH2 (EZH2-HA) and Myc-tagged NF90 (NF90-Myc) in 293T cells. Co-immunoprecipitation (Co-IP) of EZH2 revealed strong interactions between exogenously expressed EZH2 and NF90 (**Fig.1A**). Further, interactions between endogenous EZH2 and NF90 were also observed in EZH2 Co-IP assay with LNCaP and LNCaP-abl (an androgen ablation-derivative of LNCaP) cells (**Fig.1B**). Serving as a positive control, EZH2 interacted with the PRC2 core subunit SUZ12 as previously reported (**Fig.1B**). This NF90-EZH2 interaction was further confirmed by reciprocal NF90 Co-IP assays with LNCaP and LNCaP-abl cells (**Fig.1C**). NF45, a known binding partner of NF90, was included as a positive control (**Fig.1C**). Collectively, these results demonstrated strong interactions between NF90 and EZH2 under both endogenous and exogenous expression conditions, supporting NF90 as a bona fide noncanonical EZH2-associated cofactor.

**Figure 1.**
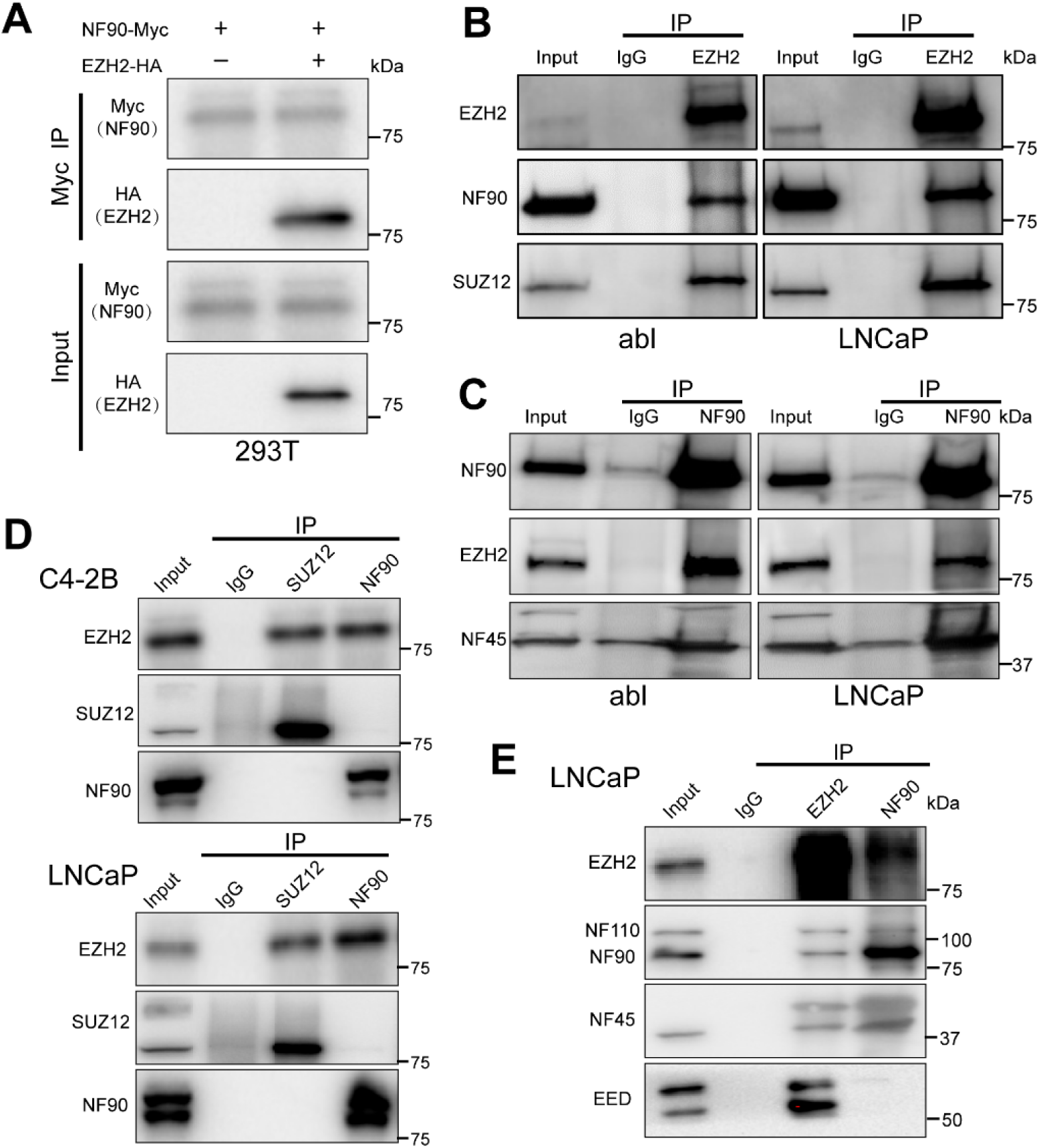
The NF90/NF110-NF45 complex interacts with EZH2 independently of PRC2. A. 293T cells overexpressing Myc-tagged NF90 with or without HA-tagged EZH2 were subjected to Co-IP assay using anti-Myc antibody, followed by western blot with indicated antibodies. B-C. LNCaP and LNCaP abl cells were subjected to Co-IP assays using anti-EZH2 (B) or anti-NF90 (C), followed by western blot analyses with the indicated antibodies. Rabbit IgG was used as a negative control. D. C4-2B and LNCaP cells were subjected to Co-IP assays with anti-SUZ12, anti-NF90, or Rabbit IgG as a negative control, followed by western blot with the indicated antibodies. E. LNCaP cells were subjected to Co-IP assays with anti-EZH2, anti-NF90, or Rabbit IgG as a negative control, followed by western blot with the indicated antibodies.

To further assess whether NF90 interacts with EZH2 through PRC2, we performed endogenous Co-IP of NF90 and SUZ12 in C4-2B and LNCaP cells. As expected, SUZ12 Co-IP assays showed robust association with EZH2 (**Fig.1D**). Surprisingly, we found that SUZ12 was completely unable to interact with NF90 protein (**Fig.1D**). Accordingly, reciprocal NF90 Co-IP assays validated that NF90 interacted with EZH2 but not with other PRC2 subunits, such as SUZ12 (**Fig.1D**). Given that NF45 is a structural partner that forms a stable heterodimeric complex with NF90/NF110 proteins and helps to stabilize them^51^, we next investigated whether EZH2 also interacts with the entire complex. Interestingly, EZH2 Co-IP assays in LNCaP cells revealed that EZH2 interacts not only with NF90 but also with NF45 and NF110 (**Fig.1E**). On the other hand, NF90 Co-IP assay showed that NF90 interacts with NF45, NF110 and EZH2, but not EED (**Fig.1E**). Taken together, our data indicate that NF90/NF110-NF45 complex specifically interacts with solo-EZH2, which exists outside of the PRC2 complex and has been shown to have polycomb-independent functions in gene activation^24^

### NF90 and EZH2 interact through their RNA-binding domains in an RNA-dependent manner

To map the EZH2 domains that interact with NF90, we co-transfected HA-tagged full-length EZH2 or a series of truncated mutants together with full-length Myc-tagged NF90 into 293T cells **(Fig.2A)**. Co-IP assays showed that all EZH2 variants, except ΔN2, were efficiently pulled down by full-length NF90, indicating that the N2 domain of EZH2 is required for this interaction **(Fig.2B).** Notably, the N2 deletion (amino acids 330-387) overlaps with the reported ncRNA-binding domain of EZH2 (residues 342-368)^52^, hence, termed as the <u>ncRNA</u> <u>domain</u> hereafter. In contrast, the enzymatically inactive EZH2 (H689A and ΔC1) mutants remained capable of interacting with NF90. The dependence on ncRNA domain implies that the NF90-EZH2 interaction may be mediated by RNA.

**Figure 2.**
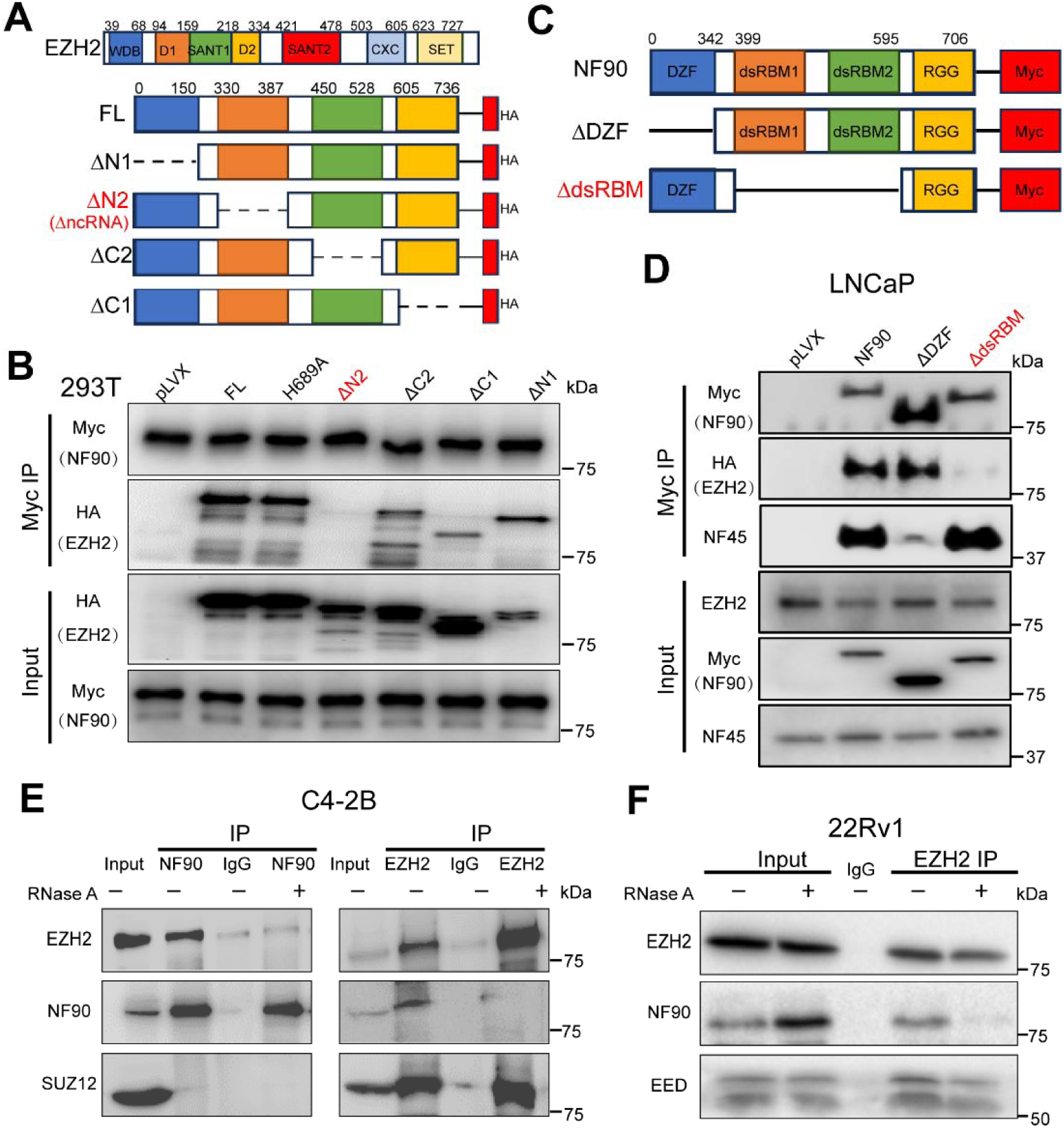
NF90 interacts with EZH2 through their RNA-binding domains. A. Domain organization of EZH2 and its truncation mutants. B. 293T cells overexpressing Myc-tagged NF90 with HA-tagged full-length EZH2 or truncation mutants were lysed and subjected to Co-IP assays with anti-Myc antibody, followed by western blot with the indicated antibodies. C. Domain organization of NF90 and its truncation mutants D. LNCaP cells overexpressing Myc-tagged full-length NF90 or its truncation mutants were lysed and subjected to Co-IP assays with anti-Myc antibody, followed by western blot with the indicated antibodies. E. C4-2B cells were lysed, treated with or without RNase A (10 μg/mL), subjected to Co-IP assays with anti-EZH2, anti-NF90, or rabbit IgG as a negative control, followed by western blot with the indicated antibodies. F. 22Rv1 cells were lysed, treated with or without RNase A (10 μg/mL), subjected to Co-IP assays with anti-EZH2 or rabbit IgG as a negative control, followed by western blot with the indicated antibodies.

Next, we mapped the NF90 domains responsible for EZH2 interaction. NF90 contains two major functional domains: DZF domain, essential for heterodimerization with NF45 to enhance protein stability and activity, and dsRBM domain, required for double-stranded RNA binding and the regulation of RNA translation and stability. We co-transfected Myc-tagged full-length NF90 or truncated mutants, ΔDZF and ΔdsRBM, together with HA-tagged full-length EZH2 into LNCaP cells **(Fig.2C)**. Co-IP assays revealed that full-length NF90 and NF90 ΔDZF, but not NF90 ΔdsRBM, efficiently pulled down EZH2, indicating that the dsRBM domain is required for EZH2 interaction **(Fig.2D)**. As expected, the NF90 ΔDZF failed to co-precipitate NF45 **(Fig.2D)**. Collectively, these results demonstrate that the interaction between EZH2 and NF90 depends on their respective RNA-binding domains, suggesting that RNA may play a critical role in mediating this interaction.

To further confirm this RNA dependency, we performed endogenous Co-IP of NF90 and EZH2 in C4-2B cell lysate with or without RNase A treatment (**Fig.2E**). The data revealed that NF90 IP enriched for EZH2, whereas EZH2 IP strongly enriched for NF90, both of which were abolished by RNase A pre-treatment (**Fig.2E**), supporting the RNA dependence of this interaction. As a control, the interaction between EZH2 and SUZ12 remained unaffected by RNase A treatment (**Fig.2E**). This RNA dependency in the NF90-EZH2 interaction was further confirmed in another PCa cell line, 22RV1 **(Fig.2F)**. Endogenous Co-IP of EZH2 in 22Rv1 cell lysate showed a strong enrichment of NF90, which was abolished by RNase A treatment **(Fig. 2F)**. Collectively, these findings support that the interaction between NF90 and EZH2 is dependent on RNA.

### NF90 cooperates with EZH2 to induce AR gene transcription

We have previously reported a critical PRC2-independent function of EZH2 in activating AR gene transcription^24^. Since NF90 is a transcriptional activator that interacts with solo-EZH2, we reasoned that it might collaborate with EZH2 in inducing the AR gene. To test this, we performed RT-qPCR analysis of AR in LNCaP cells following either EZH2 or NF90 KD. Critically, depletion of NF90 markedly reduced the mRNA levels of AR, like that caused by EZH2 KD (**Fig.3A**). This AR decrease is further supported by the reduction of TMPRSS2 and PSA, genes that are known to be directly induced by AR (**Fig.3A**). Further, RT-qPCR analyses of AR in LNCaP cells treated with actinomycin D, an inhibitor of active transcription, revealed comparable AR mRNA stability over a time course in control, EZH2 or NF90 KD cells, ruling out post-transcriptional regulation and suggesting that NF90 and EZH2 induces AR gene transcription (**Fig.S1A**). In agreement with the mRNA changes, western blot (WB) analysis confirmed that KD of either NF90 or EZH2 also decreased protein levels of AR and PSA **(Fig.3B).** To examine the effect of NF90 on AR signaling globally, we performed RNA-seq analyses of control and NF90-KD LNCaP cells. Gene set enrichment analysis (GSEA) revealed that AR-induced genes were significantly downregulated upon NF90 KD **(Fig.3C)**, whereas AR-repressed genes were upregulated upon NF90 KD **(Fig.S1B)**. This role of NF90 in activating AR transcription was further confirmed in another PCa cell line, C4-2B **(Fig.3D-E)**. RT–qPCR and western blot analyses revealed that KD of either EZH2 or NF90 in C4-2B cells markedly reduced the expression of AR and its target genes, PSA and TMPRSS2, at both the mRNA and protein levels **(Fig.3D-E)**. Conversely, overexpression of EZH2 or NF90 increased AR and its target genes expression at both RNA and protein level **(Fig.3F-G)**. Taken together, our results demonstrated that NF90, like solo EZH2, increases AR gene transcription, thereby enhancing AR signaling in PCa.

**Figure 3.**
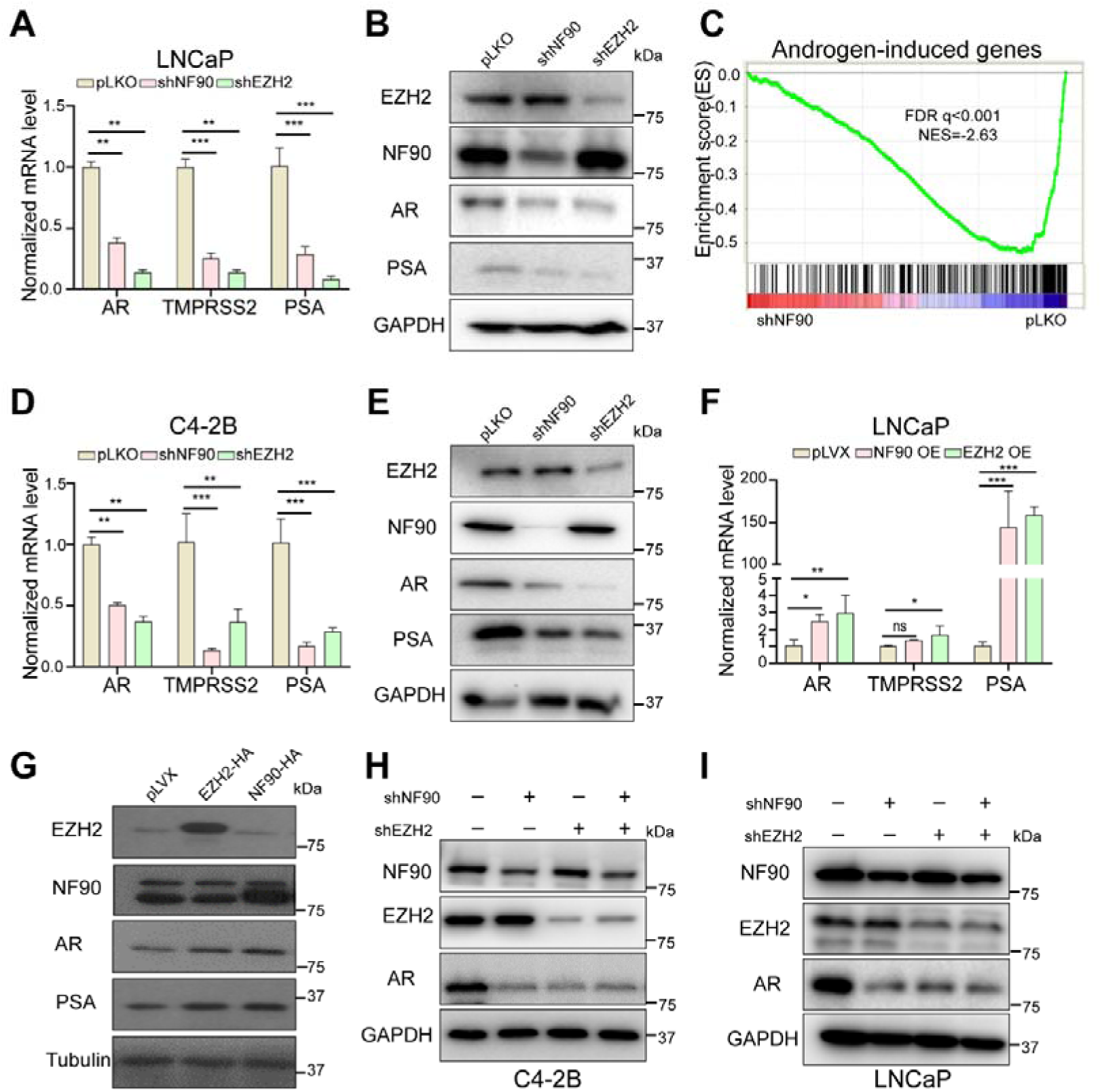
NF90 cooperates with EZH2 to induce AR transcription. A. RT-qPCR analysis of AR and its target genes, TMPRSS2 and PSA, in NF90- or EZH2-KD LNCaP cells. The y-axis shows signals after normalization to GAPDH (mean ± SEM, n=3). The statistical test is based on a one-way ANOVA paired with Dunnett’s multiple-comparison test. *, **, and *** denote P < 0.05, 0.01 and 0.001, respectively. B. Western blot analysis of AR and PSA protein levels in NF90- or EZH2-KD LNCaP cells. GAPDH was used as an internal control. C. Gene set enrichment analysis (GSEA) shows that AR-induced genes are strongly enriched in pLKO LNCaP cells (right) compared to NF90-KD LNCaP cells (left). D. RT-qPCR analysis of AR and its target genes, TMPRSS2 and PSA, in NF90- or EZH2-KD C4-2B cells. The y-axis shows signals after normalization to GAPDH (mean ± SEM, n=3). The statistical test is based on a one-way ANOVA paired with Dunnett’s multiple-comparison test. *, **, and *** denote P < 0.05, 0.01 and 0.001, respectively. E. Western blot analysis of AR and PSA protein levels in NF90- or EZH2-KD C4-2B cells. GAPDH was used as an internal control. F. RT-qPCR analysis of AR and its target genes, TMPRSS2 and PSA, in LNCaP cells overexpressing NF90 or EZH2. The y-axis shows signals after normalization to GAPDH (mean ± SEM, n=3). The statistical test is based on a one-way ANOVA paired with Dunnett’s multiple-comparison test. *, **, and *** denote P < 0.05, 0.01 and 0.001, respectively. NS denotes not significant. G. Western blot analysis of AR and PSA protein levels in LNCaP cells overexpressing NF90 or EZH2. Tubulin was used as an internal control. H-I. Western blot analysis of AR protein levels in EZH2- and NF90-KD individually or combined in C4-2B (H) and LNCaP (I) cells. GAPDH was used as an internal control.

To gain insight into how NF90 and EZH2 induce AR gene transcription, we asked whether both NF90 and EZH2 are required for AR expression and whether there are additive effects. To address this, we investigated AR expression in C4-2B cells with depletion of either EZH2 or NF90, or both. Importantly, western blot analyses revealed that the depletion of either NF90 or EZH2 alone is sufficient to abolish AR expression, and depletion of both did not produce an additive effect **(Fig.3H)**. Similar effects were also observed in LNCaP cells (**Fig.3I**), suggesting that the presence of both NF90 and EZH2 is required for AR expression. Of note, depletion of NF90 or EZH2 individually did not alter the protein level of the other, ruling out sequential events leading to AR regulation **(Fig.3H-I)**. Collectively, these results demonstrate that NF90 and EZH2 cooperatively promote AR gene transcription through an interdependent mechanism.

### NF90 and EZH2 recruit each other to the AR promoter for transactivation

We have previously shown that EZH2 directly binds to the AR promoter to induce its transcription^24^. To examine how this may be affected by NF90, we performed EZH2 ChIP-qPCR in EZH2- or NF90-KD LNCaP cells in the presence of RNase inhibitor to protect EZH2-NF90 interaction (**Fig.4A**). Interestingly, we detected a very strong EZH2 binding site at +3.3 kb downstream of the AR Transcription Start Site (TSS) (AR +3.3kb) that was abolished by EZH2 KD (**Fig.4A**). Significantly, depletion of NF90 also reduced EZH2 occupancy at this AR regulatory element, suggesting that NF90 enhances EZH2 binding. Moreover, active histone mark H3K27ac is also highly enriched at the AR +3.3 kb region (**Fig.4B**), while repressive histone mark H3K27me3 was absent (**Fig.S2A**), indicating a potential activating role of EZH2. Moreover, H3K27ac levels were remarkably reduced upon EZH2 or NF90 KD, indicating that NF90 and EZH2 bind to this regulatory element to activate AR transcription (**Fig.4B**). To further investigate whether interaction with NF90 is critical for EZH2 binding to the AR gene, we performed EZH2 ChIP-qPCR in LNCaP cells with EZH2 KD followed by rescue using wildtype (WT) or ΔncRNA-EZH2, which is unable to interact with NF90. Importantly, the results showed that ΔncRNA-EZH2 failed to bind to the AR regulatory element (**Fig.4C**). Furthermore, immunoblot analyses showed that EZH2 WT, but not ΔncRNA-EZH2, rescued AR expression in EZH2-KD cells (**Fig.4D**). Altogether, these data demonstrate that NF90 interacts with EZH2 and recruits EZH2 to the AR promoter for transcriptional activation.

**Figure 4.**
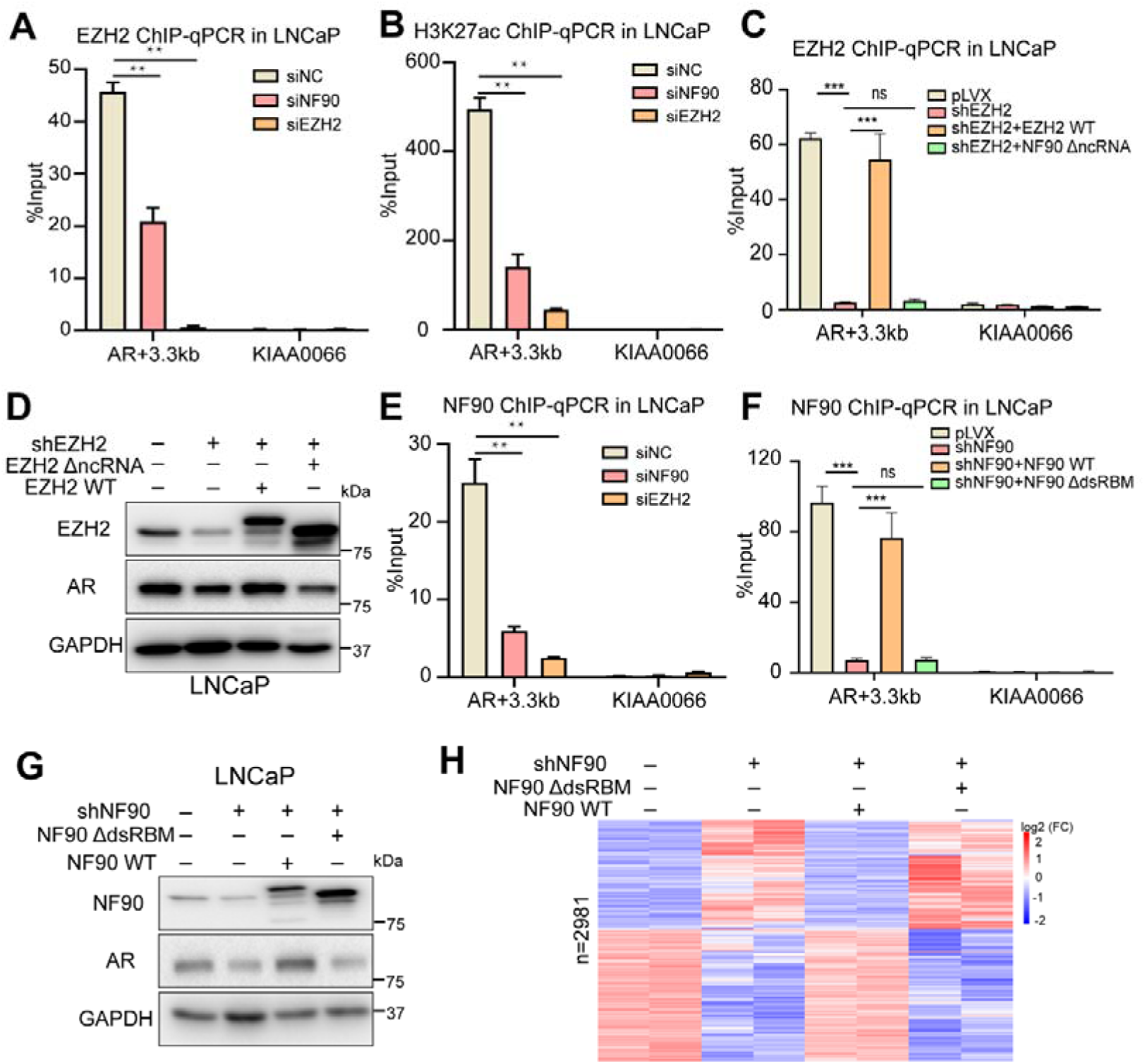
NF90 and EZH2 recruit each other to the AR promoter for transactivation. A-B. ChIP-qPCR analysis of EZH2 (A), and H3K27ac (B) at AR +3.3 kb loci in NF90- or EZH2-KD LNCaP cells. Y axis shows signals after normalization to 2% of input DNA (mean ± SEM, n=3). The statistical test is based on a one-way ANOVA paired with Dunnett’s multiple-comparison test. *, **, and *** denote P < 0.05, 0.01 and 0.001, respectively. C. ChIP-qPCR analysis of EZH2 in control and EZH2-KD LNCaP cells overexpressing control, wild-type, or ΔncRNA mutant EZH2. Data is normalized to 2% of input DNA (mean ± SEM, n=3). The statistical test is based on a one-way ANOVA paired with Dunnett’s multiple-comparison test. *, **, and *** denote P < 0.05, 0.01 and 0.001, respectively. D. Western blot analysis of AR and EZH2 protein levels in EZH2-KD LNCaP cells overexpressing either wild-type EZH2 or the EZH2 ΔncRNA mutant. E. ChIP-qPCR analysis of NF90 at AR +3.3 kb loci in NF90- or EZH2-KD LNCaP cells. The y-axis shows signals after normalization to 2% of the input DNA (mean ± SEM, n=3). The statistical test is based on a one-way ANOVA paired with Dunnett’s multiple-comparison test. *, **, and *** denote P < 0.05, 0.01 and 0.001, respectively. F. ChIP-qPCR of NF90 in control and NF90-KD LNCaP cells overexpressing control, wild-type NF90, or ΔdsRBM mutant NF90. Data is normalized to 2% of input DNA (mean ± SEM, n=3). The statistical test is based on a one-way ANOVA paired with Dunnett’s multiple-comparison test. *, **, and *** denote P < 0.05, 0.01 and 0.001, respectively. G. Western blot analysis of AR and NF90 protein levels in NF90-KD LNCaP cells overexpressing either wild-type NF90 or NF90 ΔdsRBM mutant. H. RNA-seq analysis of NF90-KD C4-2B cells overexpressing either wild-type NF90 or NF90 ΔdsRBM mutant. Heatmap shows differentially expressed genes identified (padj < 0.05) among all the groups.

To determine whether EZH2 affects NF90 binding to this AR regulatory element, we likewise performed NF90 ChIP-qPCR in LNCaP cells, and observed strong NF90 enrichment at the AR +3.3 kb site, which is abolished upon NF90 KD **(Fig.4E)**. Interestingly, depletion of EZH2 also significantly reduced NF90 binding **(Fig.4E),** indicating that EZH2 also enhances NF90 recruitment to this locus. To further investigate this, we conducted NF90 ChIP-qPCR in LNCaP cells subjected to NF90 KD followed by rescue with WT NF90 or ΔdsRMB NF90 mutant that is incapable of interacting with EZH2. Significantly, while NF90 WT formed a strong occupancy at the AR regulatory element, ΔdsRMB NF90 completely failed to bind (**Fig.4F**), suggesting that interaction with EZH2 is essential for NF90 recruitment to the AR promoter. In addition, immunoblot revealed that ΔdsRMB NF90 also failed to rescue AR expression in NF90-KD cells, different from WT NF90 (**Fig.4G**). Therefore, NF90 requires interaction with EZH2 to form stable binding at the AR promoter and activate AR transcription.

To validate these findings in additional PCa models and examine global effects, we generated NF90-KD C4-2B cells rescued with either WT NF90 or ΔdsRMB NF90 and performed RNA-seq. Bioinformatics analysis identified 2,981 genes that were differentially expressed upon NF90 knockdown, including both up- and down-regulated genes. Notably, the expression changes induced by NF90 knockdown were fully rescued by wild-type NF90, but not by the ΔdsRBM mutant (**Fig.4H**). This result suggests that the dsRBM domain of NF90 is essential not only for its regulation of AR gene but also important for its broader regulatory functions.

### NF90 induces cell growth through interacting with EZH2 and inducing AR expression

While the role of EZH2 in PCa has been well characterized, much less is known about NF90. To identify NF90 downstream genes and pathways, we performed RNA-seq analyses of C4-2B cells with or without NF90 KD. Volcano plots analysis identified 1147 down-regulated and 1234 up-regulated genes upon NF90 KD (|Log_2_ fold Change|>1 and adj. p<0.05) **(Fig.5A)**. Hallmark and KEGG enrichment analysis revealed that NF90-induced genes were highly enriched for genes involved in androgen response, being consistent with its induction of AR (**Fig.5B**). However, the most enriched genes were E2F/MYC targets and G2M checkpoint genes and the most enriched cellular process related to the cell cycle (**Fig.5B and S3A**). Interestingly, EZH2 is also a major driver of PCa cell cycle and cell proliferation, and EZH2 KD has been shown to lead to cell-cycle arrest in the G2/M phase^5,12,53^. NF90-repressed genes, on the other hand, were enriched in gene sets related to TNF-α signaling, P53 signaling pathway, and apoptosis **(Fig.5C and S3B)**. Collectively, these data suggest that NF90 is a critical regulator of the cell cycle and is likely to play important functions in promoting PCa cell growth.

**Figure 5.**
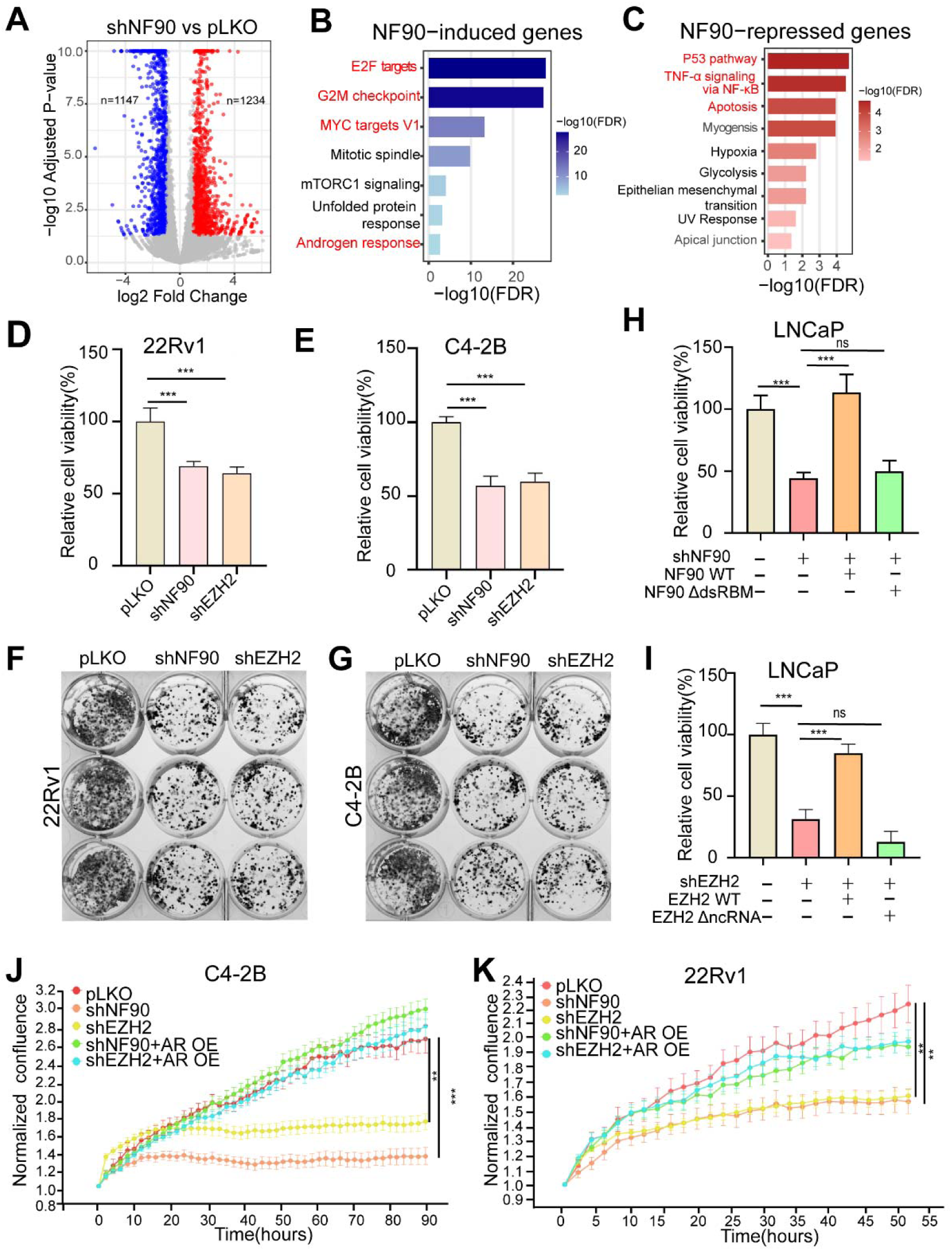
NF90 induces cell growth through interacting with EZH2 and inducing AR expression. A. Volcano plot shows differentially expressed genes (DEGs) based on RNA-seq profiles of NF90-KD C4-2B cells compared to control. DEGs that were up- or down-regulated (padjL<L0.05 and |log_2_FC|L>L1) upon NF90 KD are shown in red and blue, respectively. B-C. Hallmark pathway enrichment analysis of genes induced (B) or repressed (C) by NF90 in C4-2B cells. The X-axis shows the enrichment significance (-log10 of adjusted p-values). The Y-axis shows enriched Hallmark gene sets. D-E. Cell viability assays of 22Rv1 (D) and C4-2B (E) cells with NF90 or EZH2 KD. Cells were seeded at 7,000 cells per well in 96-well plates, and cell viability was assessed after 4 days using the WST-1 assay. Data are normalized to control cells expressing pLKO vector (mean ± SEM, n=3). The statistical test is based on a one-way ANOVA paired with Dunnett’s multiple-comparison test. *, **, and *** denote P < 0.05, 0.01 and 0.001, respectively. F-G. Representative images of Colony formation assays of 22Rv1 (F) and C4-2B (G) cells with NF90 or EZH2 KD. Cells were seeded at 5,000 per well in 12-well plates, and colonies were stained after 10 days using the crystal violet assay. H. Cell viability assays of NF90-KD LNCaP cells overexpressing control, wild-type NF90, or NF90 ΔdsRBM mutant. Cells were seeded at 7,000 cells per well in 96-well plates, and cell viability was assessed after 4 days using the WST-1 assay. Data are normalized to control cells expressing the pLKO vector (mean ± SEM, n=3). The statistical test is based on a one-way ANOVA paired with Dunnett’s multiple-comparison test. *, **, and *** denote P < 0.05, 0.01 and 0.001, respectively. NS denotes not significant. I. Cell viability assays of EZH2-KD LNCaP cells overexpressing control, wild-type EZH2, or EZH2 ΔncRNA mutant. Cells were seeded at 7,000 cells per well in 96-well plates, and cell viability was assessed after 4 days using the WST-1 assay. Data are normalized to control cells expressing pLKO vector (mean ± SEM, n=3). The statistical test is based on a one-way ANOVA paired with Dunnett’s multiple-comparison test. *, **, and *** denote P < 0.05, 0.01 and 0.001, respectively. NS denotes not significant. J-K. Cell proliferation assays of NF90- or EZH2-KD C4-2B (J) or 22Rv1 (K) cells, with or without AR re-expression. Cells were seeded at 7,000 per well in 96-well plates, and live-cell imaging was performed every 2 hours using the Incucyte system, with cell confluence quantified over time using the Incucyte software.

To delineate potential oncogenic roles of NF90 in PCa cells, we first performed cell viability assays in 22Rv1 and C4-2B cells subjected to NF90 or EZH2 KD. We observed about 40-50% reduction in live cells after four days of NF90 KD, an effect that is very comparable with EZH2 KD (**Fig.5D-E**). To examine the long-term effects of NF90 depletion on the ability of cells to survive, proliferate, and form cell colonies, we next conducted colony formation assays over a period of ten days. Importantly, depletion of NF90 remarkably diminished 22Rv1 and C4-2B colony formation to an extent similar to EZH2 KD (**Fig.5F-G**). These results suggest that NF90, like EZH2, critically regulates both short- and long-term PCa cell survival. To evaluate whether the effects of NF90 and EZH2 on cell growth are interrelated, we performed cell viability assays in LNCaP cells with NF90 KD, followed by rescue with NF90 mutants. Significantly, while WT NF90 fully restored NF90-KD LNCaP cell viability, ΔdsRBM-NF90, which is unable to interact with EZH2, completely failed to rescue cell growth (**Fig.5H**). Analogously, ΔncRNA EZH2, which has a defect in interacting with NF90, failed to restore EZH2-KD cell viability (**Fig.5I**). These results suggest that the interaction between EZH2 and NF90 is important for their ability to regulate cell growth. We thus hypothesized that their oncogenic functions may be, at least in part, mediated by their ability to induce AR expression. To test this, we used an orthogonal approach to monitor live cell growth over a long period of time with the IncuCyte system. Significantly, our data showed that depletion of NF90 or EZH2 abolished PCa cell growth and that re-expression of ectopic AR fully rescued the growth of NF90-KD or EZH2-KD cells (**Fig.5J-K**). Altogether, these results support the notion that NF90 and EZH2 induce cell growth, at least in part, through mutual interactions to induce AR expression.

### NF90 is upregulated in advanced prostate cancer, associated with EZH2 and cell cycle gene expression

To examine the clinical relevance of our discoveries, we analyzed gene expression in the TCGA prostate adenocarcinoma (PRAD) dataset. We found that the mRNA levels of ILF3, the gene encoding NF90, were significantly elevated in tumor compared to normal prostate tissues **(Fig.6A)**. Further stratification of the tumor samples revealed a stepwise increase in *ILF3* expression with rising Gleason scores, supporting a positive association with tumor aggressiveness and suggesting that *ILF3* upregulation may serve as a potential biomarker of PCa progression **(Fig.6B)**. Indeed, Kaplan-Meier survival analyses demonstrated that patients with higher levels of ILF3 showed significantly shorter biochemical recurrence-free survival (BCR) **(Fig.6C**). To further validate these findings at the protein levels, we performed immunohistochemical (IHC) analysis of NF90 on tissue microarrays (TMA) comprising 15 cases of prostatic intraepithelial neoplasia (PIN) and 36 cases of prostate cancer tissues with Gleason Grades (GG) of 3 (n=20) or GG4 (n=16). We observed that NF90 IHC staining exhibits prominent nuclear localization (**Fig.6D**), which is consistent with the nuclear distribution of EZH2. This overlapping subcellular localization suggests that NF90 and EZH2 may be in close proximity within the nucleus, providing a spatial basis for their potential interaction and coordinated regulation of nuclear processes. Importantly, our data confirmed that NF90 is significantly up-regulated in GG3 prostate cancer compared to PIN lesions and is further drastically increased in GG4 aggressive PCa **(Fig.6E**). These results support that NF90 is up-regulated in PCa and associated with more aggressive disease.

**Figure 6.**
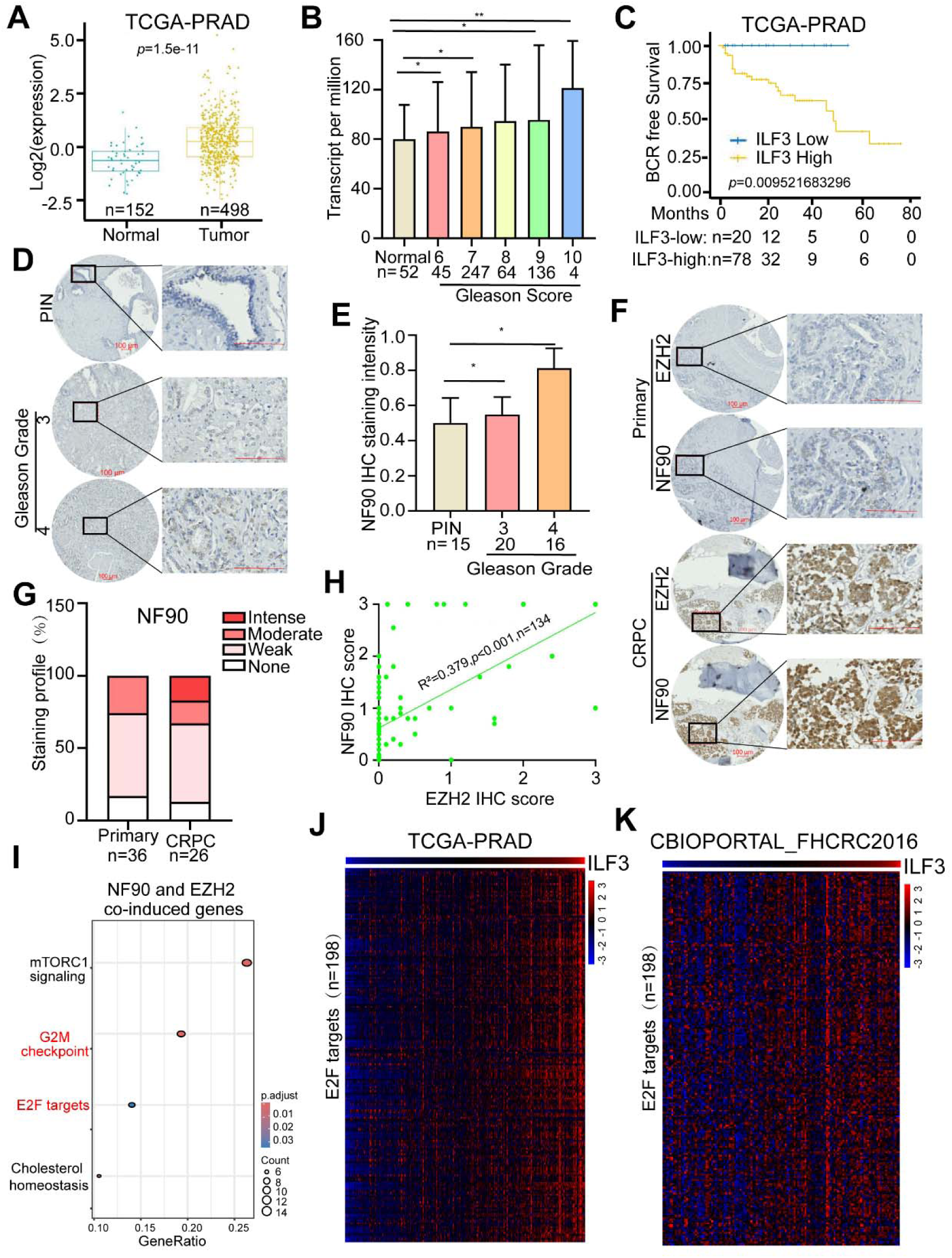
NF90 is upregulated in advanced prostate cancer, associated with EZH2 and cell cycle gene expression. A. ILF3 mRNA expression levels were compared between normal and PCa tissues in the TCGA-PRAD dataset. B. ILF3 mRNA expression levels were compared across PCa with different Gleason Scores in the TCGA-PRAD dataset. C. Kaplan-Meier analyses showing different biochemical recurrence-free survival separated by high or low ILF3 expression, using data from the TCGA-PRAD dataset. D. Representative images of IHC staining for NF90 in prostatic intraepithelial neoplasia (PIN) and PCa tissues with Gleason Grades of 3 or 4, analyzed using the primary PCa (PCF201021) TMA. The primary PCa (PCF201021) TMA comprises a total of 51 patients, including 15 PIN cases, 20 Gleason grade 3, and 16 Gleason grade 4. Scale bars:100 μm. E. Quantification of NF90 immunohistochemical (IHC) staining intensity in PIN lesions and primary PCa tissues with Gleason grades 3 or 4. The Y-axis indicates the relative IHC staining intensity. Statistical test is based on one-way ANOVA paired with Dunnett’s multiple comparison test. *P < 0.05. F. Representative images of IHC staining for NF90 and EZH2 in primary prostate tumors and CRPC samples using CRPC (UWTMA92D), comprising 26 cases in triplicate. Scale bars: 100 μm. G. Quantification of IHC staining intensities for NF90 in human primary prostate tumors and CRPC samples. The primary PCa TMAs comprise 36 patients with primary prostate cancer, including 20 with Gleason grade 3 and 16 with Gleason grade 4. The UWTMA92 TMA comprises 26 patients. The Y-axis shows the percentage of tumors exhibiting none, weak, moderate, or intense IHC staining. H. Correlation between NF90 and EZH2 IHC staining intensities in PCa TMAs. Each dot represents one tissue core on the TMA, with a total of 134 cores derived from 15 PIN, 36 primary, and 26 CRPC patients (some in replicate cores). P < 0.001 by linear regression. I. Hallmark pathway enrichment analysis of genes co-induced by ILF3 and EZH2 in LNCaP cells. The X-axis shows the ratio of differential genes in each pathway. The Y-axis lists the enriched signaling pathways. J. Heatmap showing the expression profiles of E2F target genes in PCa ranked by ILF3 expression, based on the TCGA-PRAD dataset. K. Heatmap showing the expression profiles of E2F target genes in CRPC ranked by ILF3 expression, based on the CBIOPORTAL-FHCRC2016 datasets.

To further characterize the expression pattern of NF90 in CRPC tumors, we performed IHC staining on PCa TMAs that contained triplicate cores of visceral and bone metastases (78 cores) collected from 26 patients. The data from this CRPC TMA were compared with those from the Primary TMA described above. Notably, NF90 in general showed much stronger staining in the CRPC tissues, compared to primary PCa **(Fig.6F**). In primary tumors (n=36), most cases exhibited weak to moderate NF90 staining, with none showing strong intensity. In contrast, CRPC tumors (n=26) showed a clear shift toward higher NF90 expression, with more cases displaying moderate to strong staining and fewer cases showing no staining **(Fig.6G**). These findings indicate that NF90 expression increases during the transition to CRPC. Given the potential interplay between NF90 and EZH2 in PCa, we next examined EZH2 expression in matched TMAs. Interestingly, in primary tumors, the majority of samples (approximately 80-90%) lacked detectable EZH2 staining, with only a small fraction showing weak or moderate intensity and virtually none showing strong staining (**Fig.6F, S4A**). In contrast, CRPC samples demonstrated a clear increase in EZH2 expression, with fewer cases showing no staining and more cases exhibiting weak, moderate, or strong staining (**Fig.6F, S4A**). Moreover, the IHC staining intensities of EZH2 and NF90 showed a significant positive correlation (**Fig.6H**). At the transcript levels, we also found a significant positive correlation between ILF3 and EZH2 mRNA expression levels in public PCa expression profiling datasets **(Fig.S4B)**. Additionally, western blot analysis found that NF90 protein levels are highly correlated with EZH2 protein levels across a panel of PCa cell lines **(Fig.S4C)**. Together, these results indicate that both NF90 and EZH2 are upregulated during CRPC progression, suggesting their coordinated involvement in tumor progression.

To investigate the main downstream genes and pathways of NF90 and EZH2 in PCa, we performed RNA-seq analysis in LNCaP cells following EZH2 or NF90 knockdown. Importantly, we identified 138 genes that were co-induced by EZH2 and NF90, based on the criteria of padj < 0.05 and |Log_2_ fold Change|>1 **(Fig.S4D)**. To investigate the functional pathways of these co-regulated genes, we performed Hallmark enrichment analysis, which revealed significant enrichment in cell cycle-related pathways, including G2/M checkpoint and E2F targets **(Fig.6I)**, indicating that NF90 and EZH2 coordinately promote cell cycle. To further confirm their regulation of cell cycle in human PCa, we analyzed the expression of E2F target genes in the TCGA-PRAD dataset patients. Heatmap analysis revealed that E2F target genes were markedly upregulated in tumors with high ILF3 mRNA expression **(Fig.6J).** Interestingly, a similar pattern was observed for EZH2, where tumors with elevated EZH2 mRNA levels also showed pronounced upregulation of E2F target genes, suggesting a convergent role for ILF3 and EZH2 in promoting E2F-driven transcriptional programs **(Fig.S4E).** Consistent results were also observed in CRPC samples, where both ILF3 and EZH2 showed a similar association with the elevation of E2F target genes in the CBIOPORTAL-FHCRC2016 dataset **(Fig. 6K, S4F)**. Together, these findings support a role for ILF3 in promoting PCa progression, at least in part, by enhancing cell cycle. Surprisingly, androgen-induced gene sets were not substantially upregulated in patients with high ILF3 mRNA expression, which is likely due to AR being regulated by many factors, such as gene amplification in CRPC. Collectively, these results demonstrate that ILF3 and EZH2 are coordinately upregulated in aggressive PCa and associated with cell cycle-related gene expression.

## Discussion

Although EZH2 has been extensively studied as the catalytic subunit of PRC2 that mediates epigenetic silencing through H3K27me3, accumulating evidence indicates that EZH2 also exerts PRC2-independent functions in transcriptional activation^9,23^. We have previously reported that AR is a major target of EZH2-mediated transcriptional activation in PCa, accounting for its resistance to catalytic EZH2 inhibitors^24^. Here, we demonstrate that EZH2 interaction with RNA-binding protein NF90 is essential for AR transactivation. Our Co-IP assays, combined with domain mapping, revealed that NF90 requires its dsRBM domain, whereas EZH2 requires its ncRNA-binding motif for this interaction. Interestingly, previous studies have reported that EZH2 exhibits the strongest affinity for lncRNAs among PRC2 subunits^54^, and such interaction occurs despite the absence of canonical RNA-binding motifs such as RGG or serine-rich domains, and instead relies on a 96-residue (aa340-438) intrinsically disordered region (IDR) that facilitates lncRNAs binding and recruits the entire PRC2 complex for target gene silencing^55^. Importantly, our study revealed that the RNA-binding ability of this IDR can be exploited to mediate the transactivation function of solo EZH2. Interestingly, a study has previously suggested that the N-terminal partially disordered transactivation domain (TAD, aa135-200) of EZH2 may be induced in some cancers to undergo structural transitions that enable subsequent transcriptional coactivator binding^56^. Further, several studies have reported that EZH2 interacts with coactivators such as p300 and c-Myc through this partially disordered TAD^23,56^. However, RNA and RNA-binding partners have not been shown to interact with the TAD or mediate EZH2 transactivation, indicating a function different from the RNA-binding IDR we report here. Together, our findings uncover a previously unrecognized mechanism of transcriptional regulation in which RNA functions as a molecular linker to facilitate protein-protein interactions, mediating the transactivation function of EZH2 (**Fig.7**). This mechanism adds a new dimension to the noncanonical role of EZH2 in transactivation and highlights the broader principle that structured RNA can orchestrate chromatin regulatory complexes to drive oncogenic transcription.

**Figure 7.**
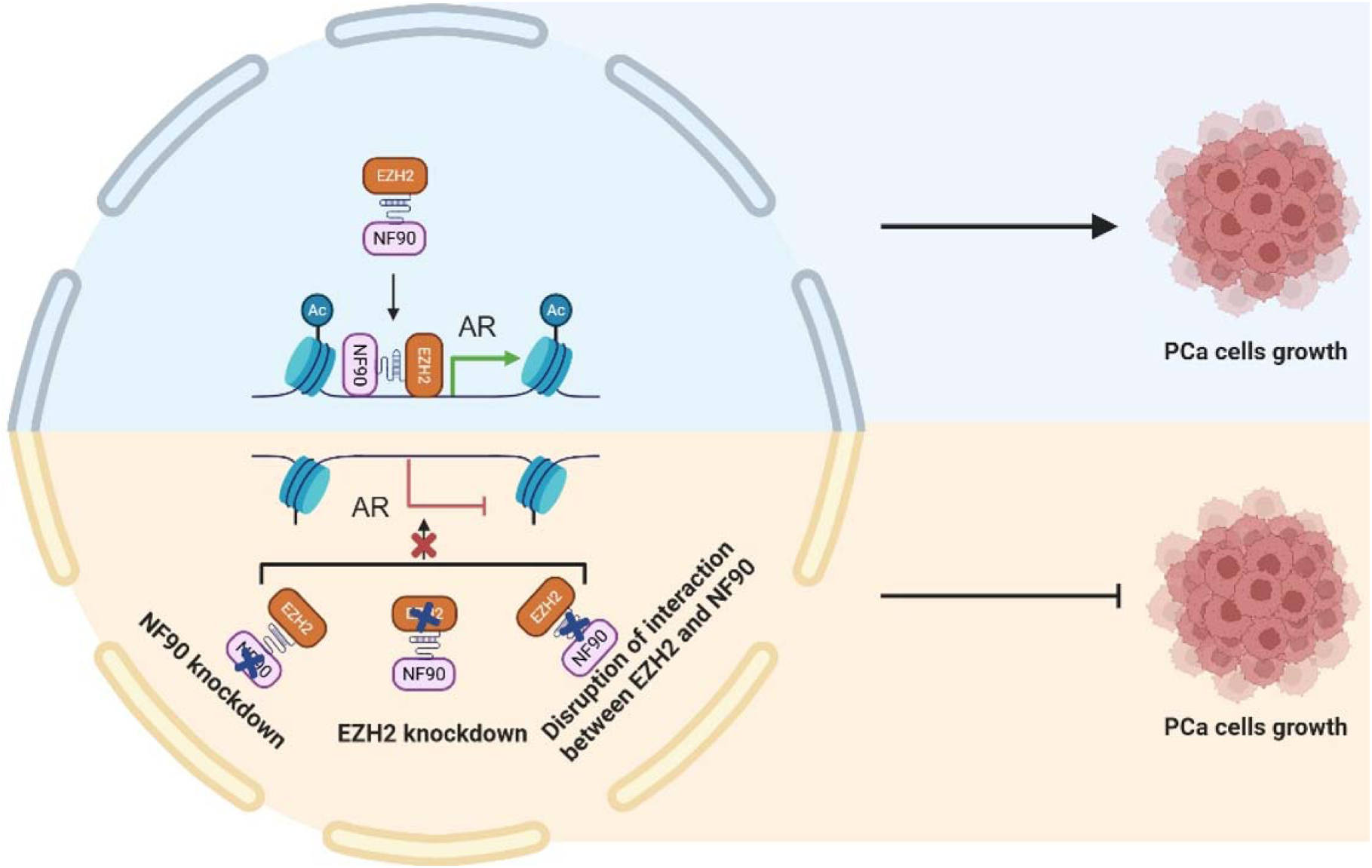
A model depicts mutual recruitment of the RNA-mediawted NF90-EZH2 complex to the AR locus, leading to AR transcriptional activation and driving PCa growth.

Our study identifies NF90 as a critical interactor of EZH2 that functions cooperatively to promote AR transcription and PCa growth. Moreover, RNA-seq analysis revealed that NF90 depletion broadly affects gene transcription, leading to downregulation of genes involved in E2F/MYC targets, G2/M checkpoint, cell cycle, and androgen response, while upregulating genes associated with apoptosis, the p53 pathway, and inflammatory signaling. These results suggest that NF90 is a master regulator of the cell cycle, similar to EZH2, which is downstream of the pRB-E2F pathway, critically mediating cell cycle^57^. We found that the ΔdsRBM mutant of NF90, unable to interact with EZH2, has an impaired ability to induce cell growth, indicating that NF90 might collaborate with EZH2 in regulating the cell cycle. However, it should be noted that the dsRBM of NF90 is important for a plethora of NF90 activities and target genes. Interestingly, a recent study has reported that AR cooperates with E2F1 to induce novel transcription networks that promote PCa^58^. Further, EZH2 has been shown to enhance E2F1 activity in a methylation-independent manner^59,60^. These studies indicate multiple potential mechanisms through which NF90 may promote cell-cycle progression and warrant further investigation, which, however, falls outside the scope of the current work.

Our analysis of clinical mRNA datasets and TMA of patient samples showed that high NF90/ILF3 expression is associated with higher Gleason scores and shorter biochemical recurrence-free survival, suggesting a potential prognostic role. NF90 is upregulated in aggressive PCa and CRPC and positively correlates with EZH2 and cell-cycle gene expression. Accordingly, like EZH2, NF90 strongly promotes cell proliferation and colony formation, and this effect is at least in part dependent on AR expression. These results establish NF90 as a promising therapeutic target in advanced PCa. However, the cooperative function of NF90 and EZH2 in inducing AR not only has major impacts on PCa growth but might also raise important implications for lineage plasticity. While EZH2 has long been a major therapeutic target in PCa, recent evidence suggests that EZH2 suppression, surprisingly, leads to lineage plasticity and the transition of PCa to AR-low or AR-null neuroendocrine prostate cancer (NEPC), a highly aggressive and therapy-resistant lineage state^61^. This double-edged sword effect of EZH2 may, at least in part, be mediated by AR, the loss of which is a critical event in NEPC progression. Hence, the NF90-EZH2 axis not only elucidates how EZH2 supports AR transcription in adenocarcinoma but also offers a conceptual framework for understanding how disruption of this regulatory module could potentially drive AR loss and foster lineage plasticity. Future studies investigating strategies to target NF90 and how its suppression influences PCa across different stages of disease progression will be essential.

There are several limitations in our study. We focus on delineating the important roles of RNA and RNA-binding protein NF90 in mediating EZH2 transcriptional activation, utilizing AR as a prototype gene. We did not examine the global chromatin targets of these proteins, as it is beyond the scope of the current study. We demonstrated that the NF90-EZH2 interaction is essential for their binding to the AR promoter, but we did not investigate the mechanisms underlying their recruitment to chromatin, which will be an important focus for future studies. In addition, this study did not pinpoint an lncRNA that is essential for NF90 and EZH2 interaction, due to the promiscuous RNA-binding effects of EZH2, suggesting a lack of selectivity ^62^. Lastly, given the striking effects of NF90 on regulating the cell cycle and cell growth in vitro, we feel an animal experiment is unnecessary, in line with recent NIH guidelines to reduce animal use.

## Materials and Methods

### Cell lines, chemical reagents, and antibodies

The PCa cell lines LNCaP, 22Rv1, and C4-2B, as well as the human embryonic kidney cell line HEK293T, were obtained from the American Type Culture Collection (ATCC, Manassas, VA). LNCaP, 22Rv1 and C4-2B were cultured in RPMI 1640 medium supplemented with 10% fetal bovine serum (FBS) and 1% penicillin and streptomycin. HEK293T cells were maintained in Dulbecco’s modified Eagle’s medium (DMEM) with 10% fetal bovine serum (FBS) and 1% penicillin and streptomycin. All cell lines were authenticated within six months of culture and routinely tested for mycoplasma contamination. All antibodies used in this study are listed in Supplementary Table 1.

### Plasmids and Lentivirus Infection

The shRNAs targeting NF90 and EZH2 were inserted into the pLKO.1-TRC lentiviral vector (Addgene, 10878). Primers and oligonucleotides used in this study are listed in Supplementary Table 2. All plasmids were verified by sequencing. Stable overexpression and KD were reached by lentiviral infection. HEK293T cells were transfected with psPAX2, pMD2.G, and the target gene at a ratio of 2:1:1. Lentiviral supernatant was collected 48 hours posttransfection and filtered through a 0.45 μm membrane. Lentiviruses supplemented with 8 μg/mL polybrene were used to infect PCa cells. For most shRNA-mediated KD experiments, cells were harvested four days postinfection.

### siRNA Transfection

Cells were seeded in 6-well plates at 50-70% confluency the day before transfection. SiILF3 or siEZH2 targeting the gene of interest or a non-targeting control siRNA (scramble) was transfected using Lipofectamine RNAiMAX (Thermo Fisher Scientific, Waltham, MA) according to the manufacturer’ s instructions. Briefly, siILF3 or siEZH2 was diluted in Opti-MEM (Gibco) and mixed with the transfection reagent, followed by incubation at room temperature for 5-20 minutes to allow complex formation. The siRNA-reagent complexes were then added to the cells, and cells were cultured at 37°C with 5% CO₂. Knockdown efficiency was assessed 72 hours post-transfection by qRT-PCR and/or Western blot analysis.

### Co-immunoprecipitation (Co-IP)

Cells were resuspended in IP lysis buffer (50mM Tris-HCl, pH7.4, 150mM NaCl, 1mM EDTA, 1% Triton X-100, 1□×□Roche protease inhibitor cocktail) and incubated on ice for 20□minutes. The lysates were then centrifuged at 14,000 rpm for 20 minutes at 4□°C, and the supernatant was transferred to pre-chilled 1.7 mL microcentrifuge tubes. Save 30 μL of the lysate as the input control, mix it with 10 μL of 4× SDS loading buffer, and boil at 95□°C for 10 minutes to denature proteins. Store the input samples at -80□°C until use. Add 2 μg of the desired antibody to the remaining lysate and incubate overnight at 4□°C with gentle rotation. Vortex the protein A/G beads thoroughly to resuspend them, then take 25 μL of beads per sample and wash them at least once with 1 mL of IP wash buffer or PBS. Add 25 μL of protein A or G magnetic beads to the lysate and incubate with gentle rotation at 4□°C for 2 hours. Place the Eppendorf in the magnetic rack to magnetize the beads and discard the supernatant. Wash the beads six times with 1 mL of IP wash buffer by rotating at 4□°C for 3 minutes each time. After each wash, place the tube on the magnetic rack and remove the wash buffer completely. Elute the bound proteins by adding 50 μL of 2× SDS loading buffer. Incubate at 95□°C for 10 minutes with shaking to denature the proteins. Place the tube on the magnetic rack again and transfer the supernatant (elute) to a new microcentrifuge tube. Load the eluates onto SDS-PAGE gels for WB analysis using the corresponding antibodies. All antibodies used in this study are listed in Supplementary Table 1.

### RNA extraction, RT–qPCR, and RNA-seq

RNA was extracted using the nucleospin RNA kit (Takara, San Jose, CA) according to the manufacturer’s recommended protocol. Then, 500□ng of RNA was reverse transcribed into complementary DNA (cDNA) using the ReverTra Ace qPCR RT Master Mix kit (Diagnocine, Hackensack, NJ) according to the manufacturer’s recommended protocol. qPCR was performed using 2X Universal SYBR Green Fast qPCR Mix (Abclonal, Cat#RK21203) and QuantStudio 3 (Thermo Fisher, Waltham, MA). All primers used here are listed in Supplementary Table 1. For RNA-seq, total RNA was isolated as described above. RNA-seq libraries were prepared from 0.5 μg of high-quality DNA-free RNA using NEBNext Ultra RNA Library Prep Kit, according to the manufacturer’s instructions. The libraries passing quality control (equal size distribution between 250 and 400 bp, no adapter contamination peaks, no degradation peaks) were quantified using the Library Quantification Kit from Illumina (Kapa Biosystems, KK4603). Libraries were pooled to a final concentration of 10 nM and sequenced single-end using the Illumina HiSeq 4000.

### RNA-seq data processing and analysis

RNA-seq data quality was first assessed using FastQC (v0.12.1), which confirmed high sequencing quality across all samples. Adapter trimming and low-quality base removal were performed using fastp (v1.0.1). Ribosomal RNA contamination was filtered out with SortMeRNA (v4.3.7). The cleaned reads were aligned to the human hg38 (UCSC) reference genome using STAR (v2.7.11b), yielding high mapping rates for all samples. Gene-level counts were summarized and subjected to differential expression analysis using DESeq2 (v1.46.0). Significantly differentially expressed genes (DEGs) were defined based on adjusted *P*-value (padj) and log2 fold-change thresholds. Functional enrichment of DEGs was performed using clusterProfiler (v4.14.0), revealing pathway alterations consistent with the observed transcriptional changes. Collectively, these analyses identified robust transcriptomic differences between the experimental groups.

### ChIP–qPCR

For EZH2, NF90, H3K27ac, and H3K27me3 ChIP assays, LNCaP cells were dual cross-linked using 1% formaldehyde (Thermo Fisher, Cat#28908) and DSG (disuccinimidyl glutarate). Cells were first treated with 50 mM DSG for 15 min, followed by 1% formaldehyde for 10 min at room temperature with gentle rotation, and the reaction was then quenched with 0.125 M glycine for 5 min. A total of 10 million cells were used for each EZH2 and NF90 ChIP, while 5 million cells were used for each H3K27me3 or H3K27ac ChIP. During chromatin-protein isolation, 5U/µl RNase inhibitors were included to preserve both the EZH2-NF90 interaction and the association of EZH2 with chromatin. Chromatin was sheared to an average fragment size of 200-600 bp using an E220 focused ultrasonicator (Covaris,Woburn, MA). Supernatants containing chromatin fragments were pre-cleared with protein A agarose beads (Millipore, Burlington, MA) for 40□min and incubated with a specific antibody overnight at 4□°C on a nutator (antibody information is listed in Supplementary Table 1), then 50 μL of protein A agarose beads were added and incubated for 2 h. The beads were washed twice with 1□×□dialysis buffer (2□mM EDTA, 50□mM Tris-Cl, pH 8.0) and four times with ChIP wash buffer (100□mM Tris-Cl, pH 9.0, 500□mM LiCl, 1% NP40, 1% deoxycholate). The antibody/protein/DNA complexes were eluted with elution buffer (50□mM NaHCO3, 1% SDS), the crosslinks were reversed and the DNA was purified using DNA Clean & Concentrator-5 kit (ZYMO Research, Irvine, California). Elute the DNA in 30 µl of water. If necessary, further dilute the samples 300-fold and use 2-3 µl per PCR reaction. Perform qPCR using established positive and negative control genes.

### WST-1 cell proliferation assay, Incucyte live cell imaging, and colony formation assay

Cell proliferation assay was performed with WST-1 (Clontech Laboratory, Mountain View, CA) as described by the manufacturer’s instructions. Briefly, C4-2B and 22Rv1 cells were seeded into 96-well plates, followed by the addition of 10 µl WST-1 reagent per well and a 2-h incubation. The absorbance was then measured at 450 nm using a spectrophotometer. For the Incucyte assay, C4-2B and 22Rv1 cells were counted using a Countess automated cell counter (Life Technologies, Carlsbad, CA) and seeded into 96-well plates in triplicate. Photomicrographs were acquired every 2 hours using an Incucyte live-cell imaging system (Essen Biosciences, Ann Arbor, MI). Cells were quantified using the Incucyte software (Essen Biosciences, Ann Arbor, MI).

For the colony formation assay, LNCaP and C4-2B cells (5□×□10^3^ cells per well) were seeded in 12-well plates. The cells were fixed with 4% paraformaldehyde after ten days of growth and stained with 25% methanol containing 0.5% crystal violet for 15 minutes. The colonies were imaged with ChemiDoc (BIO-RAD, Hercules, CA).

### IHC and TMA analysis

The primary PCa (PCF201021) TMA (number of patients=51, number of cores = 65) was generated by the Northwestern University Pathology Core and approved by the Northwestern University Institutional Review Board. CRPC (UWTMA92D) TMA containing metastatic CRPC specimens was obtained from the University of Washington Medical Center Prostate Cancer Donor Program, which is approved by the University of Washington Institutional Review Board (IRB). UWTMA92 Array D was investigated, consisting of 1 mm core in triplicates of visceral metastases and bone metastases (number of cores = 78) collected from 26 patients.

TMAs were constructed using 1-mm cores from primary or CRPC tissues. EZH2 and NF90 staining on the TMAs was performed using standard Immunohistochemistry (IHC) procedures with the ImmPRESS® Excel Amplified Polymer Kit (Vector Laboratories), according to the manufacturer’s instructions. Briefly, formalin-fixed paraffin-embedded (FFPE) TMA tissue sections were deparaffinized and hydrated, followed by antigen retrieval, which was accomplished by heating at 99-100□°C for 15 min in 1x citrate buffer, pH 6.0 (Sigma, C9999-1000ML). After antigen retrieval, the tissue sections were incubated with BLOXALL blocking solution to quench endogenous peroxidase activity and then were blocked with 2.5% normal horse serum. After blocking, the tissue sections were incubated with primary antibodies, followed by incubating with amplifier antibody (Goat Anti-Rabbit IgG for rabbit primary antibodies) or directly incubated with ImmPRESS polymer reagent (for mouse primary antibodies). After ImmPRESS polymer reagent incubation, the tissue sections were incubated in ImmPACT DAB EqV working solution until the desired staining intensity develop, then the tissue sections were counterstained with hematoxylin, mounted with mounting medium and imaged with the Zeiss Axioscan 7. Antibody information for IHC was provided in Supplementary Table 1. NF90 and EZH2 immunostaining were scored in a blind fashion using a score of 0 to 3 for intensities of negative, moderate, or intense and multiplied by the percentage of stained cancer cells.

### Statistical analysis

For each independent in vitro experiment, at least three technical replicates were performed. Most in vitro experiments were repeated independently three times, and few were repeated twice. Results are expressed as mean ± standard error (mean ± SEM). A two-tailed unpaired Student’s t-test was used to evaluate data consisting of 2 groups. A one-way ANOVA paired with Dun nett’s multiple comparison tests was used to evaluate data consisting of 3 and more groups. The results were considered significant at p < 0.05. Kaplan–Meier analyses of free survival and overall survival of patients were performed using the log-rank test.

## Data availability

All RNA-seq data generated in this study are prepared to be deposited in the Gene Expression Omnibus with GSE311720.

## Acknowledgments

This work was partially supported by the Pathology Core Facility and Imaging Core Facility at Emory University, as well as the Pathology Core Facility at Northwestern University. We thank Dr. Michael B. Mathews for providing the NF90/NF110 plasmids. Fundings support for the work includes the NIH/NCI grant R01CA293596, the prostate cancer SPORE P50CA180995, and the Department of Defense grant PC160856. JDL is supported by NCI P30 CA247796 and R01CA266078.

## Contributions

JY conceived the project, and JY and YW designed the experiments. YW performed most of the experiments. LP did ChIP-qPCR experiments. XL provided guidance about the project design and repeated ChIP-qPCR experiments. MZ and JCZ conducted bioinformatic and statistical analyses. SS performed routine mycoplasma detection of cell culture and repeated some Co-IP experiments. YL assisted with some experiments. HS performed IHC experiments. JA performed some Co-IP and protein detection experiments. JDL provided valuable comments on the project and edited the manuscript. XY helped with the TMA and pathology. YW, HC, and JY wrote the manuscript. All authors reviewed and approved the manuscript.

