## Supplementary information for "RNA-Binding Protein NF90 Mediates Polycomb-Independent Transactivation by EZH2 to Promote Cancer Growth"

**Supplementary Figures:**

**Figure S1.** NF90 cooperates with EZH2 to induce AR transcription

**Figure S2.** NF90 and EZH2 recruit each other to the AR distal promoters for transactivation

**Figure S3.** NF90 induces cell growth through interacting with EZH2 and inducing AR expression

**Figure S4.** NF90 is upregulated in advanced prostate cancer and positively correlated with EZH2, and cell cycle gene expression

**Figure and Figure Legend**

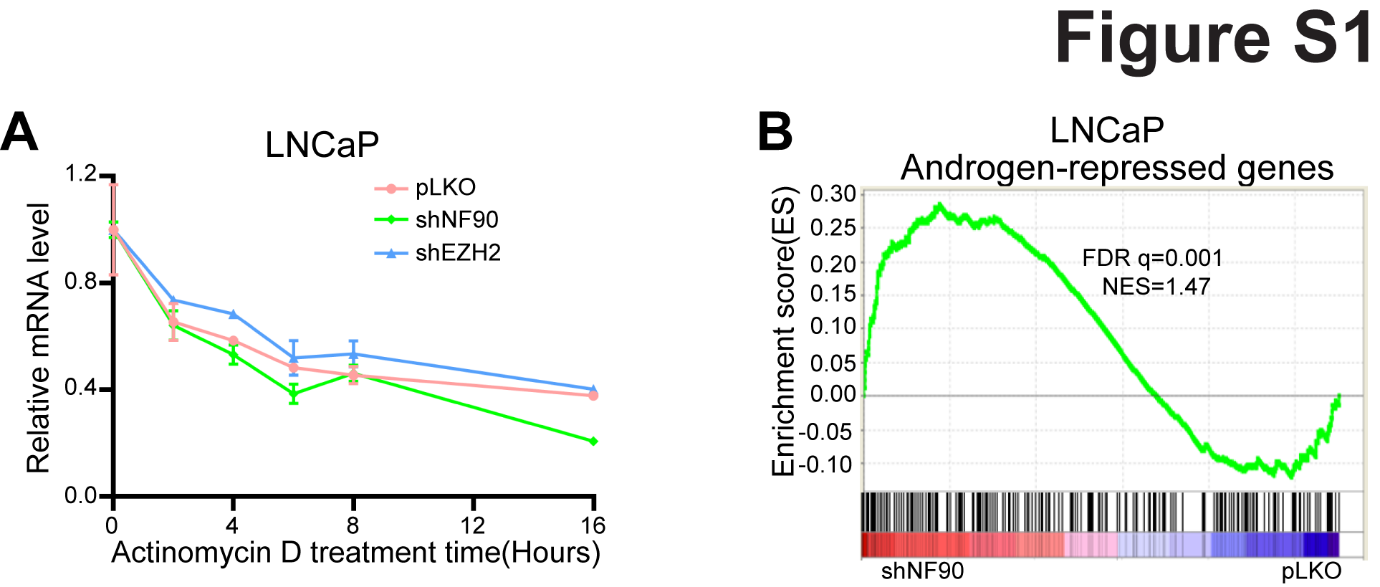

**Figure S1. NF90 cooperates with EZH2 to induce AR transcription**

A. RT-qPCR analysis of AR in EZH2 or NF90-KD LNCaP cells treated with actinomycin D (5μg/mL) for 2, 4, 6, 8, and 16 hours. Y-axis shows signals after normalization to an internal control (GAPDH) and then to the pLKO mock control.

B. GSEA analysis shows that AR-repressed genes are strongly enriched for in NF90-KD LNCaP cells compared to pLKO cells.

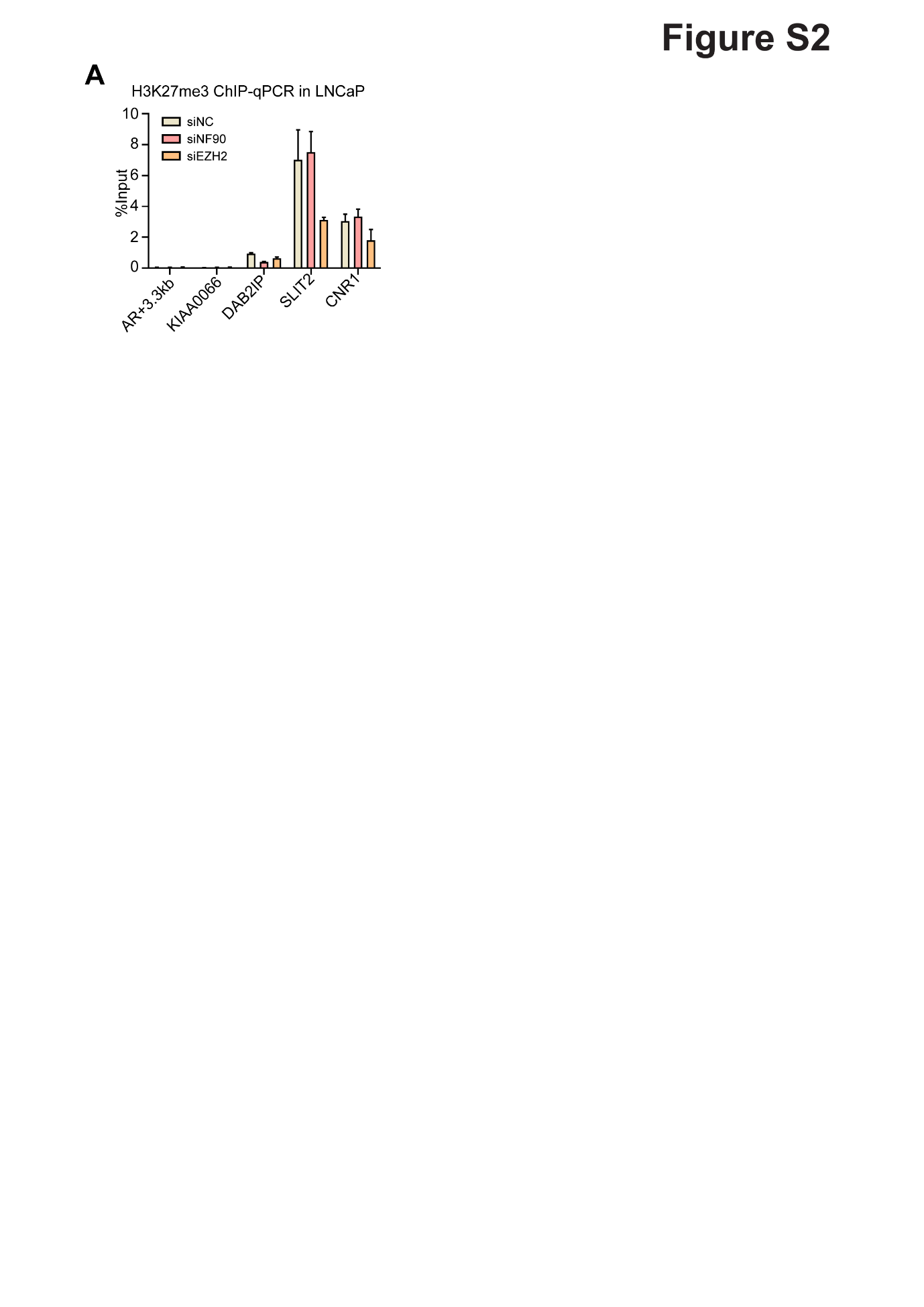

**Figure S2. NF90 and EZH2 recruit each other to the AR distal promoters for gene transactivation**

A. ChIP-qPCR analysis of H3K27me3 at AR +3.3 kb loci in NF90- or EZH2-KD LNCaP cells. Data was normalized to 2% of the input DNA (mean ± SEM, n=3). The statistical test is based on one-way ANOVA paired with Dunnett’s multiple comparison test. *, **, and *** denote P < 0.05, 0.01 and 0.001, respectively.

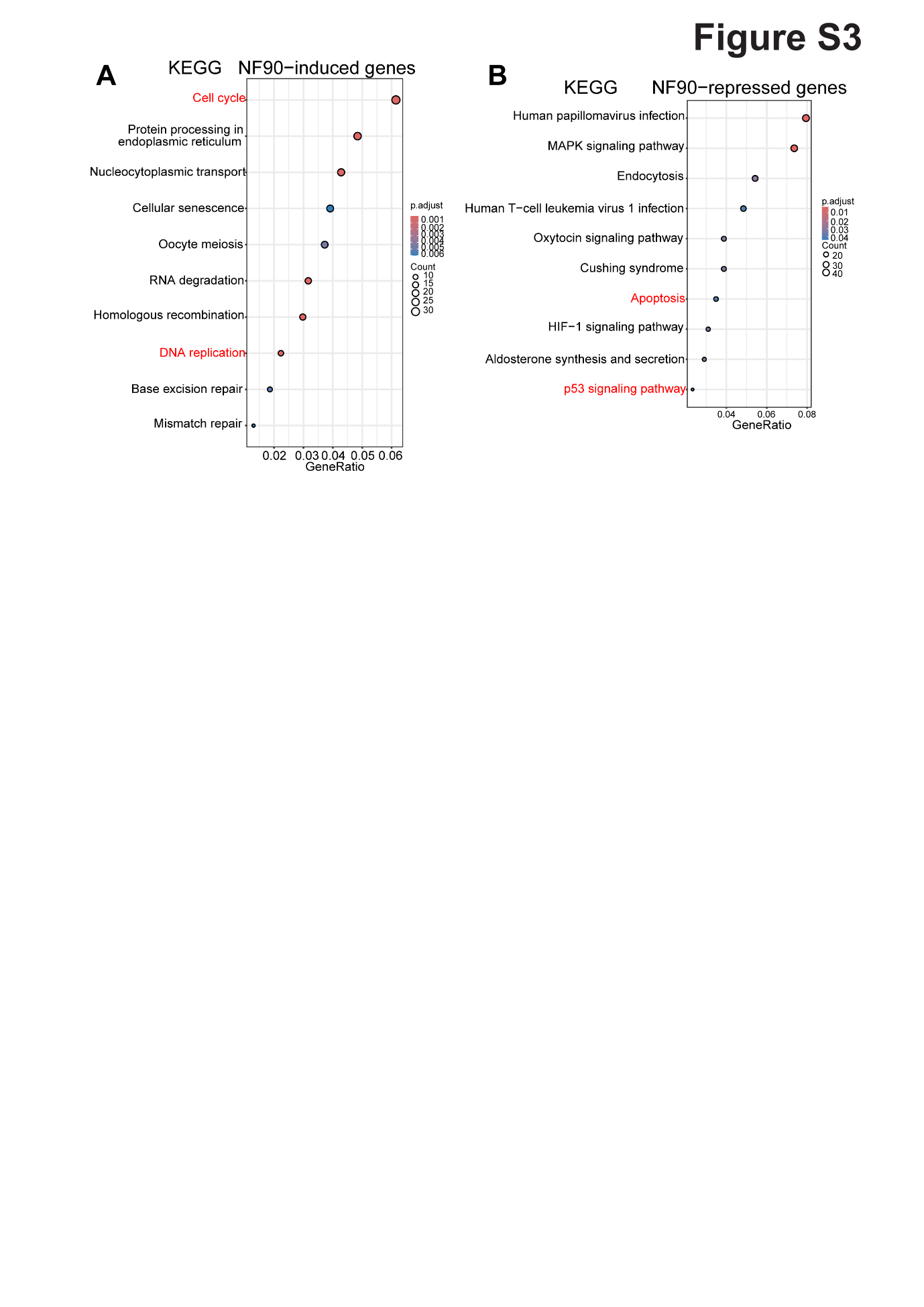

**Figure S3.NF90 induces cell growth through interacting with EZH2 and inducing AR expression**

A-B. Kyoto Encyclopedia of Genes and Genomes (KEGG) pathway analysis of NF90-induced (A) or NF90-repressed (B) genes in C4-2B cells for enrichment in different signaling pathways. The X-axis shows the ratio of differential genes in each pathway. The Y-axis lists the enriched signaling pathways.

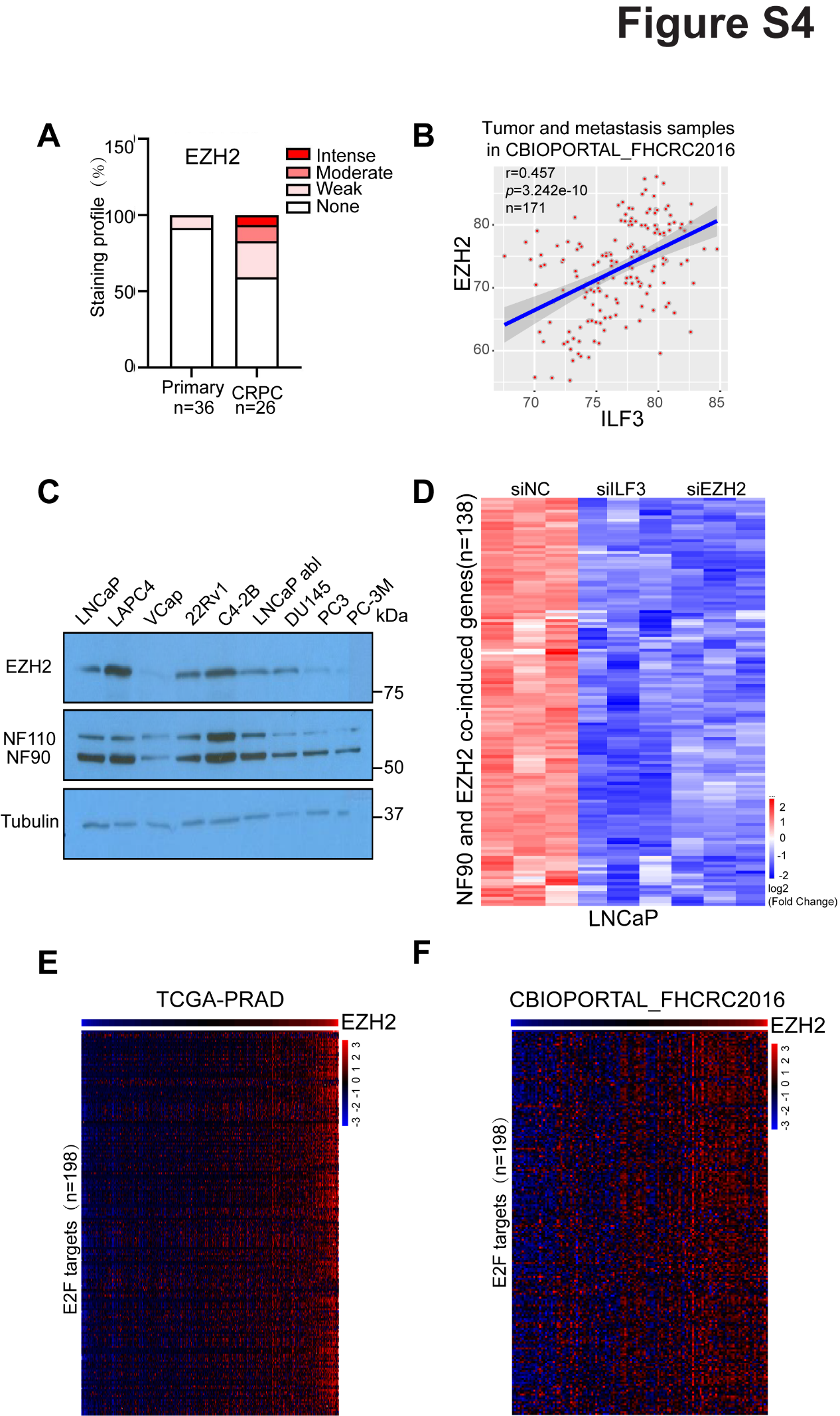

**Figure S4. NF90 is upregulated in advanced prostate cancer and positively correlated with EZH2, and cell cycle gene expression**

A. Quantification of IHC staining intensities for EZH2 in human primary prostate tumors and CRPC samples. The Y-axis shows the percentage of tumors exhibiting none, weak, moderate, or intense IHC staining.

B. Correlation between ILF3 and EZH2 in PCa using data from the CBIOPORTAL-FHCRC2016 datasets. Scatter plot showing a positive correlation between ILF3 and EZH2 mRNA expression levels in PCa samples from the CBIOPORTAL-FHCRC2016 (r = 0.457, p = 3.242e-10, n = 171).

C. Western blot analysis of the EZH2 and NF90/N110 protein levels in a panel of PCa cell lines. Tubulin is used as a loading control.

D. Heatmap showing the expression profiles of EZH2- and ILF3-induced gene sets derived from RNA-seq analysis of NF90- and EZH2-KD LNCaP cells compared with siNC.

E. Heatmap showing the expression profiles of E2F target genes in PCa ranked by EZH2 expression, based on the TCGA-PRAD dataset.

F. Heatmap showing the expression profiles of E2F target genes in PCa ranked by EZH2 expression, based on the CBIOPORTAL-FHCRC2016 datasets.
